# Scale Cortisol Signatures in Cultured Nile Tilapia (*Oreochromis niloticus*) Regulates Mitogen Activated Protein Kinase Signalling Pathway to Modulate Chronic Stress

**DOI:** 10.1101/2024.02.14.580283

**Authors:** John Gitau Mwaura, Paul Oyieng Angienda, Clabe Wekesa, Eunice Toko Namuyenga, Philip Ogutu, Patrick Okoth

## Abstract

Chronic stress poses a challenge to aquaculture, with cortisol and glucose traditionally used as stress markers. Recent doubts about the reliability of scale cortisol as a chronic stress determinant have surfaced due to its role in calcium homeostasis. While cortisol affects gene expression in stress responses, its impact on metabolic pathways in cultured Nile tilapia remains understudied. We explored the connection between cortisol signatures of chronic stress in Nile tilapia and the Mitogen Activated Protein Kinase (MAPK) signaling pathway, vital for cell processes. Juvenile Nile tilapia were exposed to varying ammonia concentrations and stocking densities, evaluating growth performance, stress levels, RNA sequencing, and differential gene expression. Results indicate a positive correlation between stressors and blood glucose, plasma cortisol, and scale cortisol concentrations. Fish in 0.8 mg/L ammonia exhibited heightened plasma glucose and cortisol, while those in 1.2 mg/L showed increased scale cortisol. Elevated stocking densities also correlated with higher stress markers. Importantly, cortisol levels rose with ammonia concentration and stocking density, negatively impacting growth. The MAPK signaling pathway, crucial for cell processes, exhibited significant downregulation with increasing ammonia concentrations, suggesting sensitivity to stress. Six genes in this pathway were significantly enriched following ammonia treatment, including Dual Specific Protein Phosphatase 1, Nuclear Hormone Receptor 38, Heat Shock Protein 1, Myelocytomatosis oncogene homologue, Growth arrest and DNA damage inducible alpha a, and Mitogen Activated Protein Kinase 4. This study contributes valuable insights for optimizing fish welfare and production by unraveling the complex relationship between chronic stress and the MAPK pathway in aquaculture.

## 1.0 Introduction

Aquaculture intensification is on the rise driven by increasing demand for fish and fisheries products coupled with the decline in capture fisheries (FAO, 2022). There is pressure to produce more fish per area in order to feed the ever-growing world population (Rodriguez-Barreto et al. 2019). Unfortunately, this pressure has often resulted in farming more fish in limited space, leading to overcrowding and overstretching of the water environments. Contamination of fish culture water with agro-chemicals such as abamectin (Mahboub et al. 2022) and oxyfluorfen (Mansour et al. 2023) have been documented as sources of stress in fish. Industrial pollutants such as Bisphenol-A (BPA) (Hamed et al. 2021) have also been found to cause stress in fish lowering their productivity. Other pollutants such as lead (Abdel-Tawwab et al. 2023) zinc (Hamed 2023; Hamed et al. 2022), silver nitrate nanoparticles (Tohamy et al. 2022) have also been implicated in causing chronic stress in fish. Fish husbandry practices can also be a source of stress. The ensuing stress can either be acute or chronic depending on the intensity and duration of exposure. The stress ensuing from crowded sub-optimal water environments lowers the productivity and compromises fish welfare (Martos-Sitcha et al. 202; Aketch et al. 2014). The increased concentration of ammonia and its derivatives in the aquaculture environment is one of the riskiest stresses for freshwater fish (Zeitoun et al. 2016). Aquaculture utilizes high protein feeds which upon being metabolized yields ammonia as the by-product (Godoy-Olmos et al. 2022). In addition, fish faecal matter and other decomposing organic matter increases ammonia concentrations in pond water. Furthermore, stressed fish are also known to release higher levels of ammonia than unstressed fish. High stocking densities and ammonia are common occurrences in aquaculture systems and can evoke a stress response. Stocking density is an important factor determining the profitability of an aquaculture venture and farmers tend to increase stocking with the aim of increasing productivity (Oke and Goosen 2019). High stocking density has been shown to produce a chronic stress situation in Gilthead sea bream (Jia et al. 2022), the same husbandry practice has also been shown to induce chronic stress which elevates the levels of plasma cortisol and glucose in *O. niloticus* (Aketch et al. 2014).

The primary stress response is the activation of the Hypothalamo-Pituitary Interrenal (HPI) axis which culminates in the release of glucocorticoids from inter renal cells located in the head kidney (Lai et al. 2021). This leads to the production of corticosteroids mainly cortisol (Kennedy and Jauz 2022; Aidos et al. 2020) which then triggers several physiological and metabolic pathways (Vercauteren et al. 2021; Sapolsky et al. 2000). The action of cortisol is mediated through the glucocorticoid receptor (GR) which is a cytosolic receptor (Cabrera-Busto et al. 2021). Cortisol has been shown to regulate the expression of genes involved in growth, metabolism and immune function ; Balasch and Tort 2019). Cortisol up-regulates pathways involved in energy substrate mobilization, including gluconeogenesis, down-regulating energy demanding pathways such as growth, reproduction and immune functions. The reaction of fish to stressors can be determined by analysing the concentration of cortisol produced especially in plasma (Kennedy and Jauz 2022; Sadoul and Geffroy 2019). Nonetheless, there is variability in the plasma cortisol content that is often seen during sampling (Samaras et al. 2021; Harper and Wolf 2009) due to the invasive nature of the sampling process. Plasma cortisol rises rapidly after initiation of a stressful event and falls back to baseline values within 24 hrs (Kennedy and Jauz 2022; Laberge et al. 2019). This makes plasma cortisol a poor indicator for chronic stress in fish. Plasma glucose is another parameter that has found a lot of application in stress determination (Odhiambo et al. 2020) but like plasma cortisol, it is a poor chronic stress indicator (Sadoul and Geffroy 2019). A good chronic stress indicator must not fluctuate due to capture, must depict the adrenal or interrenal activity and should also reflect the integrated stress hormone concentrations over a long period of time (Kennedy and Jauz 2022). The study by Laberge et al. (2019) demonstrated that scale cortisol meets the above criteria thus qualifying it as a suitable biomarker for chronic stress.

Cortisol has been found to accumulate in fish scales (Vercauteren et al. 2021; Carbajal et al. 2018; Aerts et al. 2015) making it a suitable matrix for a retrospective study of cortisol production and thus for assessment of chronic stress in fish. Several studies have been carried out on scale cortisol of wild caught fish (Vercauteren et al. 2021; Roque d’orbcastel et al. 2021) but information about scale cortisol of cultured fish remains scanty and poorly documented. Unlike other matrices such as water, serum/plasma, faeces and mucus which reflects short-term circulating cortisol levels, fish scale cortisol represents average levels acquired mainly through passive diffusion over a period of time [30]. Vercaunteren *et al*. [18] showed that accumulation of scale cortisol in common dab (*Limanda limanda*) occurs proportionally to the circulating cortisol levels. Scale cortisol determination is relatively non-invasive and does not suffer the rapid elevation witnessed during sampling as in plasma and mucus (Pickering and Pottinger 1989). However, unlike hair, feathers and wool that have been used to study chronic stress in birds and mammals, scales sometimes are involved in calcium and cortisol homeostasis (Zenth et al. 2022). This makes the scale cortisol to be less reliable as an accurate measure of HPI activity in fish unless investigated alongside other predictors of stress (Kennedy and Jauz 2022).

Cortisol regulates several pathways involved in stress management in fish. One such pathway is the (Mitogenic Activated phosphate Kinase) (MAPK) signalling pathway. MAPK signalling pathway is essential in synchronization and regularization of cell survival, motility, differentiation, and proliferation (Cargnello and Roux 2011), metabolism, and apoptosis. MAPK signalling pathway plays a central role in hypoxia adaptation in fish (Zhu et al. 2013). Saduol and Geffroy (2019) demonstrated that cortisol modulates MAPK pathway by suppressing the phosphorylation of the extracellular signal-regulated kinase (ERK1/2), p38MAPK and c-Jun N-terminal kinase/stress-activated protein kinase (JNK). This occurs via the Dual-Specificity Phosphatase –1 (DUSP-1) also known as Mitogen activated protein Kinase -1 (MPK -1) (Wang et al. 2016). DUSP-1 regulation is the key mechanism of action of glucocorticoids such as Scale cortisol and has been used as a bio-marker for chronic stress in fish (Vercauteren et al. 2021; Roque d’orbcastel et al. 2021) but information about use of a combination of two or more markers remains scanty and poorly documented. Inhibition of MAPK may be a likely means of inhibiting cell growth (Eanes and Patel 2016). MAPK signalling pathway represents a highly ubiquitous and evolutionarily conserved mechanism of eukaryotic cell regulation. The multiple MAPK pathways present in all eukaryotic cells enable coordinated and integrated responses to diverse stimuli (Kyriakis and Avruch 2012). Therefore the aim of the present study was to elucidate the use of MAPK signalling pathway and cortisol signatures in assessing the occurrence of chronic stress in cultured Nile tilapia.

### 2.0 Materials and Methods

Hand sexed male Nile tilapia juveniles mean weight 25 ±1.25 g; total length 10 ± 0.35 cm were obtained from Ilara fish hatchery in Kakamega County in Western Kenya, and reared in the laboratory for 2 weeks prior to the start of the experiment. During the acclimatization and experimental period, the fish were maintained on a 12h light and 12h dark photoperiod cycle. They were fed to satiation two times a day with commercial feed 2mm diameter pellet size containing 32% of gross protein and 3,500 Kcal/kg of digestible energy. During the feeding, the fish were observed for 30 min and any uneaten feed was siphoned out.

### 2.1 Experimental Set-up

#### 2.1.1 Ammonia Stress Experiment

A total of 525 fish were randomly divided into seven groups and stocked in 21 white circular 500 L polyethylene tanks in a static system aerated by blowers connected to air stones. Three replicates of 25 fish per tank as guided by (Odhiambo et al. 2020) were used. The tanks were fitted with calibrated thermostats set at 26°C for temperature control. The dissolved oxygen concentration and temperature were measured with a digital oximeter (Hydrolab MSIP-REM-HAH-QUANTA (USA). The dissolved oxygen was always maintained at levels above 6mg/L [28]. This was achieved by blowing in air using an aquarium air pump connected to air stones for effective diffusion while the temperature was maintained at 26 ± 1°C for the period of the experiment (Aerts et al. 2015). The pH was measured using a digital pH meter and maintained at 6.8 ± 0.2 (Roque d’orbcastel et al. 2021). The spring water used during this experiment had a natural pH of 6.8. The treatments were distributed as follows: The first (control) group were maintained in natural water without addition of ammonia throughout the growth period. Six groups of 25 fish per tank were maintained at different concentrations of unionized ammonia (0.4, 0.8, 1.2, 1.6, 2.0 and 2.4 mg/L) throughout the growth period (Pickering and Pottinger 1989). Each of the treatments above were replicated twice with the treatments being randomly assigned within the blocks.

#### 2.1.2 Stocking Density Experiment

A total of 1,185 juvenile Nile tilapias were randomly distributed in 21 white circular polyethylene tanks (500 L). Each tank was assigned a treatment as follows: The control group were stocked at 20 fish per tank and replicated twice. Six groups of fish were maintained at different stocking densities (25, 40, 55, 70, 85 and 100 fish per tank). Each of the treatments above were replicated twice and randomly assigned within the blocks. Every day, 20% water exchange was carried out to allow for the siphoning of the left-over feeds and the excrement in each tank (Zenth et al. 2022). The fish were then reared for 70 days and their growth monitored every fortnight. Growth was monitored by determining the total length, weight, condition factor, growth rate and specific growth rate (SGR) (Cargnello and Roux 2011; Zhu et al. 2013). During the growth period, the fish were fed to satiation twice per day.

### 2.2 Sample Collection

Three fish from each treatment were randomly sampled from each tank. The selected fish were anaesthetised in 3-aminobenzoic acid ethyl ester Methane-Sulfonate (MS-222, Sigma, USA) at a dosage of 100mg/L (Wang et al. 2016) for physiological parameter measurements and the scales removal for cortisol level determinations. Ten scales were removed from along the lateral line using a clean forceps. The fish were then sacrificed and 2g of muscle in the dorsal caudal region quickly removed and ground in liquid nitrogen.

### 2.3 RNA Extraction and Sequencing

The resulting powder was immediately mixed with TRIzol® reagent (Invitrogen, Waltham, MA, USA). RNA extracted from it following the manufacturer’s instructions. Briefly, the mixture was incubated for 10 min at room temperature and RNA extracted by addition of 200µL of chloroform chilled on ice for 3 min and centrifuged at for 15 min at 4 C. The upper aqueous layer was transferred to a new tube and RNA was precipitated by addition of isopropanol. The pellet was washed twice in 75% ethanol and air dried at room temperature for 15 min. The pellet was the dissolved in 50 µL RNAse free water (Thermo Fisher Scientific, Walthan, MA, USA). DNA was digested by addition of 50µL DNAse 1 (Thermo Fisher Scientific, Walthan, MA, USA). RNA was quantified by measuring absorbance at 230 nm, 260 nm and 280 nm using Nano Vue spectrophotometer (GE Healthcare, Chicago, IL, USA). The 260/280 and 260/230 ratio was used to determine the quality and only samples with both ratios greater than 2 were used for sequencing. cDNA synthesis was carried out using SuperScript IV Reverse Transcriptase kit (Thermo Fisher Scientific, Walthan, MA, USA). Second strand synthesis was carried out using NEBNext®Ultra™IIRNA library prep kit for Illumina® (New Egland Biolabs GmbH, Ipswisch, MA, USA). The mRNA library constructed was sequenced with Illumina Miseq (Illumina, Inc. US Illumina) at Novogene Company Limited (Cambridge, UK). The count normalisation and differential expression analysis was performed using DESeq2 v1.30.1 (Dong et al. 2018) in R v4.1.2. Differentially Expressed Genes (DEGs) were identified as significantly expressed if their adjusted p-value was less than 0.05 and their fold change greater than 2.0.

### 2.4 Whole Blood Glucose Determination

The sampled fish were anaesthetised by submerging in MS-222 for one minute. Whole blood was obtained by venepuncture in the tail vein. Blood glucose levels were determined using a hand-held one touch ultra-glucose meter (MD300) and test strips manufactured by TaiDoc. Technologies Corporation and supplied by MD instruments INC. according to (Hoppstädter and Ammit, 2019; Thorarensen et al. 2015; Abdel-Tawab et al. 2014). The whole blood was applied on the test strips that were fixed in the hand-held glucose meter and the glucose concentrations were read in mMol.^-1^.

### 2.5 Plasma Cortisol Determination

The sampled fish were terminally anaesthetised by quick submersion in 5ml/L of 2-Phenoxyethanol for one minute. 2 ml of blood were collected by venepuncture in the tail vein using disposable K_2_EDTA (0.5M, pH 8.0) treated syringe and gauge 30 needle. The blood was maintained on ice throughout the sampling process and span down (1000 x g at 4^0^C for 15 minutes). The plasma was then removed and transferred to a new Eppendorf tube and frozen at - 80^0^C until analysed. The plasma cortisol was assayed by competitive inhibition enzyme linked immunosorbent assay (ELISA) using a Fish Cortisol Elisa kit (Cusabio. Wuhan, China) following the manufacturers instruction. The plasma was diluted with the sample diluent (1:100) before the test. The plates were pre-coated with rabbit anti-cortisol antibody. On to each well 50 µl of either sample or the cortisol standards were aliquoted in duplicates. Two blank wells were included in the assay where no sample or standard was added. This was immediately followed by addition of 50 µl of antibody except for the blank. The plate was shaken gently and incubated for 40 minutes at 37^0^C. The plate was then washed three times with wash buffer and 100 µl of HRP-conjugate to each well except the blank. The plate was then covered with adhesive strip and incubated for 30mins at 37^0^C. The plate was then washed five times in wash buffer before adding 90 µl of substrate solution (3,3’,5,5’-tetramethylbenzidine (TMB) to each well and incubating at 37^0^C for 20 min in a dark place. The reaction was stopped by adding 50 µl of stop solution (0.5 M sulphuric acid) and absorbance measured immediately at 540 nm. A standard curve was constructed using eight standards and the concentration of the cortisol in the plasma samples was read from this standard curve (Figure 1).

**Figure 1:**
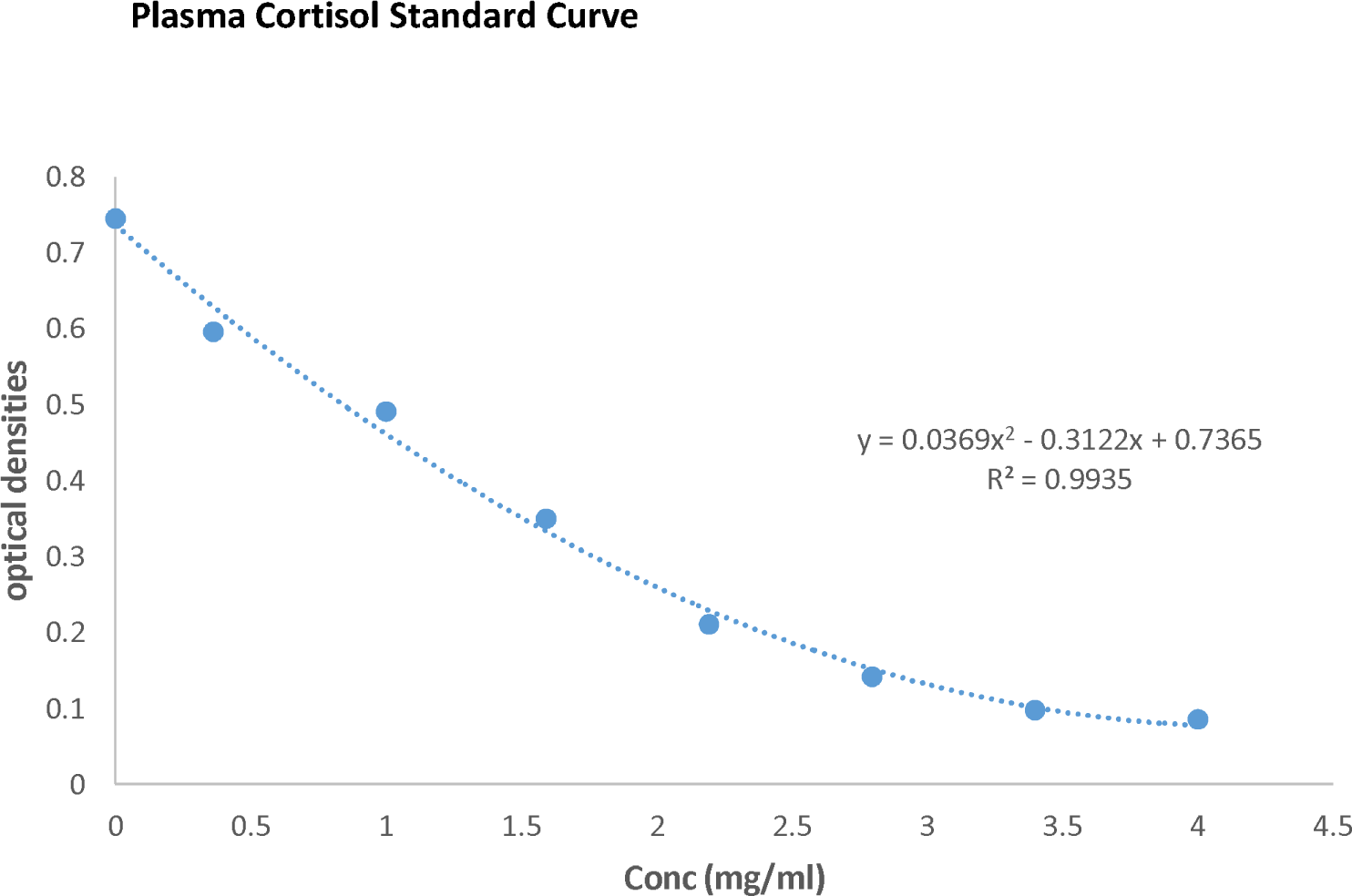
Plasma cortisol standard curve used to determine the plasma cortisol concentrations in the plasma obtained from fish in ammonia and stocking density treatments

### 2.6 Scale Cortisol Determination

Measurement of scale cortisol levels was performed using the plasma method described above with modification. After anaesthetising the fish, ten scales were removed from along the lateral line using a clean forceps (Sadoul and Geffroy 2019). The scales were washed thrice with 3 ml of isopropanol for 2.5 min for each wash and air dried on a tissue paper at room temperature. The dried scales were then chopped into a fine particle using a clean pair of scissors. The cortisol was extracted by incubating 0.100 ± 0.001 g of the fine powder in 8 ml of methanol for 16 hrs at 35^0^C. The mixture was then centrifuged at 3000 g for 10 minutes and the supernatant decanted into a fresh tube and the solvent evaporated under a stream of nitrogen at 38^0^C. The resulting powder was dissolved in 200 µl of sample diluent and cortisol level determined as in plasma cortisol determination described above. A standard curve for the determination of scale cortisol (Figure 2) was constructed as in 2.4 above.

**Figure 2:**
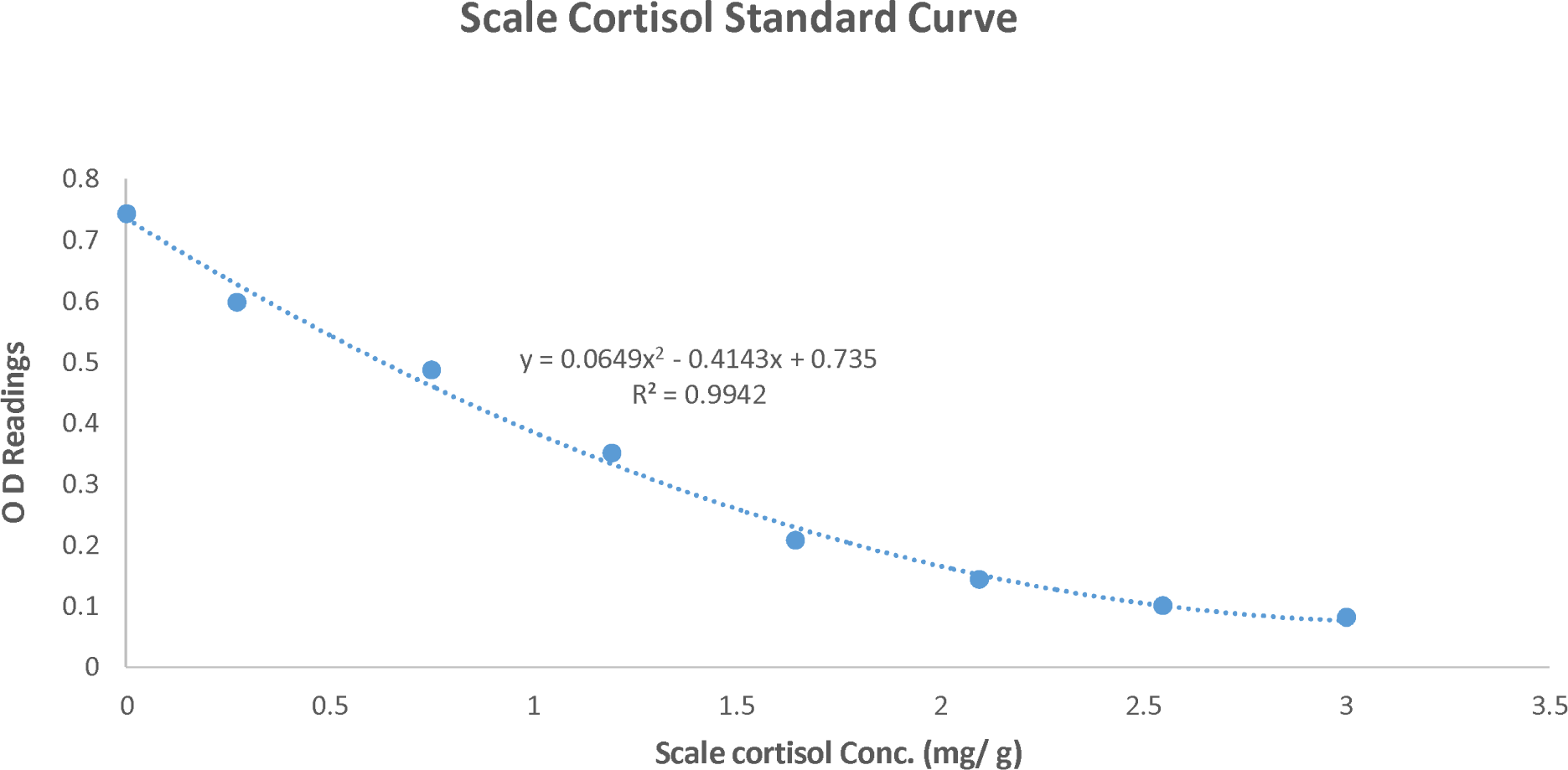
Scale cortisol standard curve used to determine the scale cortisol concentrations in scales from fish in ammonia and stocking density treatments.

### 2.7 Statistical analysis

Data were subjected to Levene’s test to test for homogeneity of variance before proceeding to perform one way ANOVA (IBM SPSS for Windows version 26) at 95% confidence interval. Levene’s tests for the two treatments gave p-values greater than 0.05.

## 3.0 Results

### 3.1 Plasma cortisol levels

The effect of chronic stress on plasma cortisol levels is as shown in tables 1 and 2 below. There was a significant difference in the mean± SD plasma cortisol concentration between the controls and the ammonia treatments (p< 0.05) i.e. 3.65 ± 0.73 and 5.11 ± 1.00 ng/ml respectively (Table 1). The results clustered into in three clusters (low, Middle and High). The mean cortisol levels increased with increase in the ammonia concentration in the culture water.

**Table 1:**
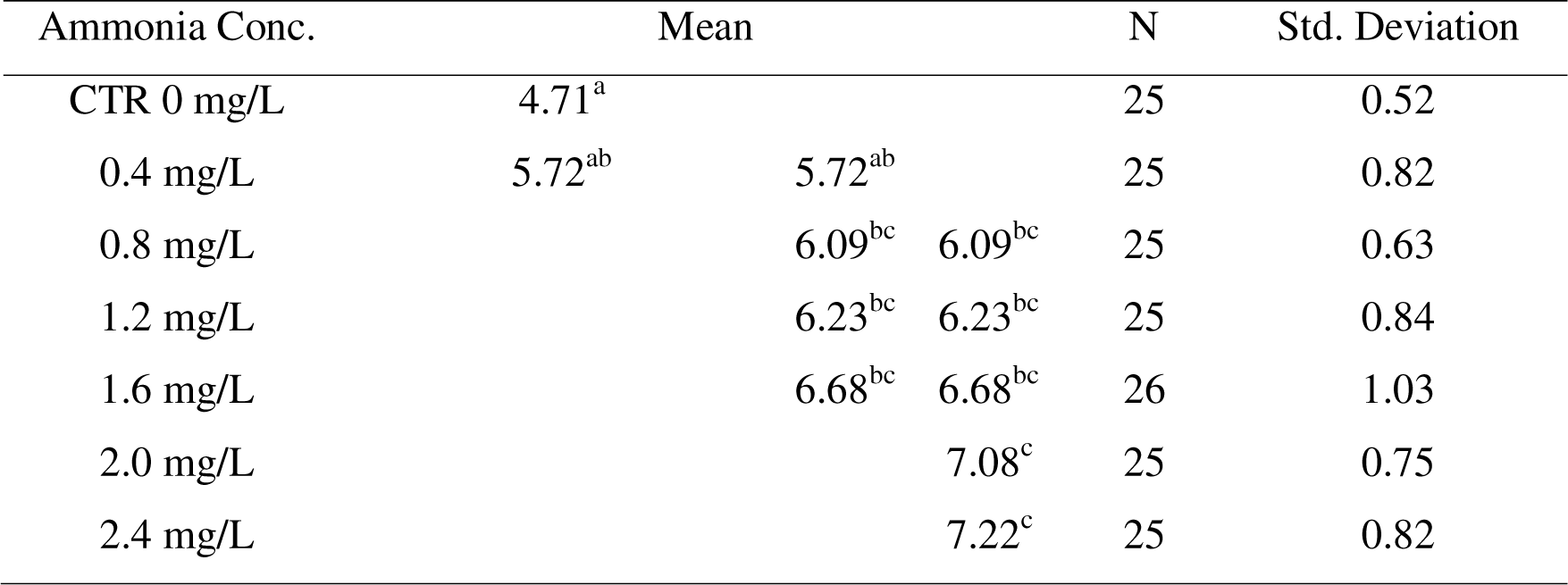
Mean plasma cortisol concentrations following ammonia treatment. The superscripts indicate significant difference at p<0.05 of the mean plasma cortisol concentration following ammonia treatment. Treatment with 0.8 mg /L and above had significantly higher plasma cortisol concentrations compared to the control.

**Table 2:**
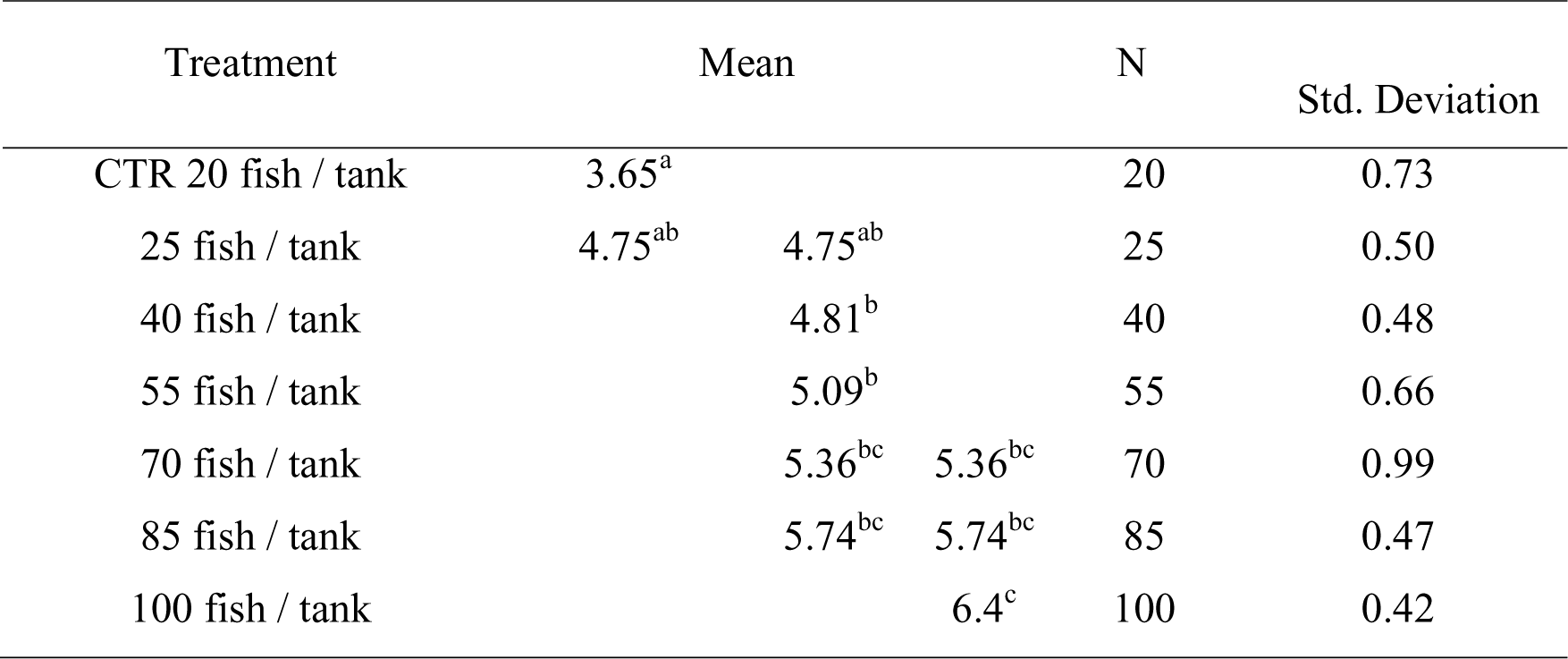
Mean plasma cortisol concentrations after subjecting to stocking density treatment. The superscripts indicate significant difference at p<0.05 of the mean plasma cortisol concentration following stocking density treatment. Treatment with 40 fish / tank and above had significantly higher plasma cortisol concentrations compared to the control.

There was a significant difference in the mean± SD plasma cortisol concentration between the controls and the stocking density treatments (p< 0.05) i.e. 3.65 ± 0.73 ng/ml and 5.11 ± 1.00 ng/ml respectively (Table 2). The result clustered into three clusters (low, Middle and High). The cortisol concentration was positively correlated to the stocking density.

### 3.2 Blood glucose levels

The effect of chronic stress on blood glucose levels is as shown in tables 3 and 4. The glucose concentration in blood was significantly higher in ammonia treatments compared to the controls (i.e., mean ± SD: 26.91±4.32 vs. 18.23±4.1 mg/dL respectively (Table 3). There was a positive correlation between the ammonia concentration and the glucose levels. Separation of means also clustered the treatments into three categories; low, medium and high.

**Table 3:**
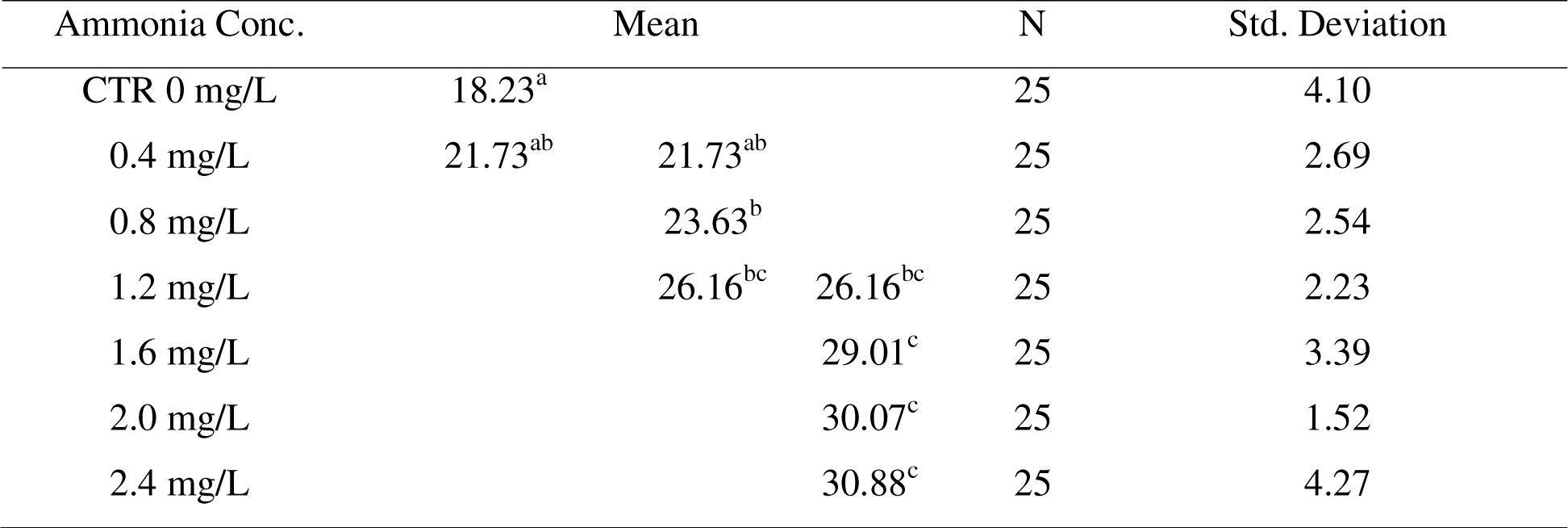
Mean plasma glucose concentration for ammonia treatment. The superscripts indicate significant difference at p<0.05 of the mean plasma glucose concentration following ammonia treatment. Treatment with ammonia concentration of 0.8 mg/L and above had significantly higher plasma glucose concentrations compared to the control.

**Table 4:**
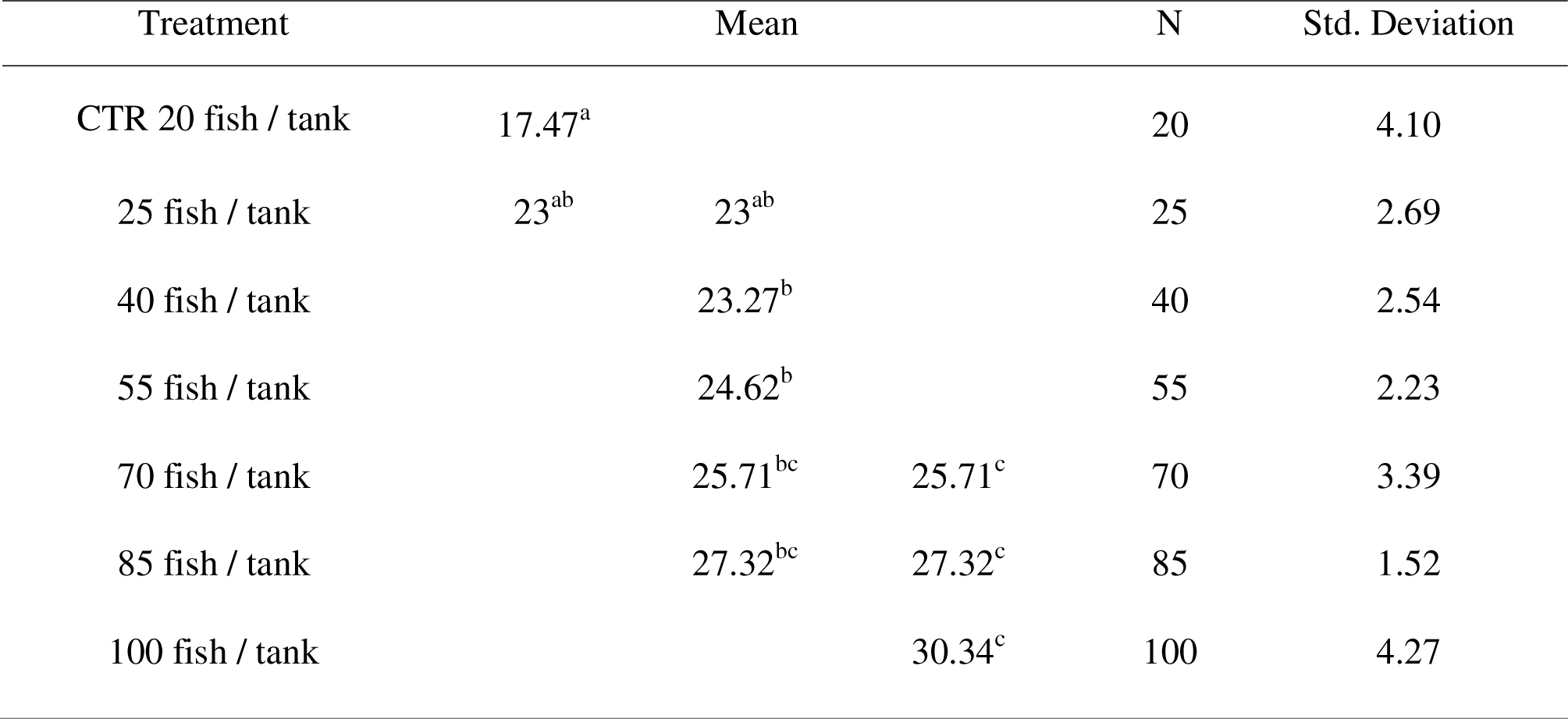
Mean blood glucose concentration for stocking density treatment. The superscripts indicate significant difference at p<0.05 of the mean plasma glucose concentration following stocking density treatment. Treatment with 40 fish / tank and above had significantly higher plasma glucose concentrations compared to the control.

Stocking density also had a profound effect on the blood glucose concentrations as shown in table 4 below. The mean glucose concentration ranged from 17.47 ± 4.10 mg/dL for the control to 30.34 ± 4.27 mg/dL for the highest stocking density used (100 fish per tank). Although there was a significant difference between the treatments when compared to the controls, there was no significant difference when the treatments were compared to each other.

### 3.3 Scale cortisol levels

The effects of chronic stress on scale cortisol concentration was as outlined in tables 5 and 6 below.

**Table 5:**
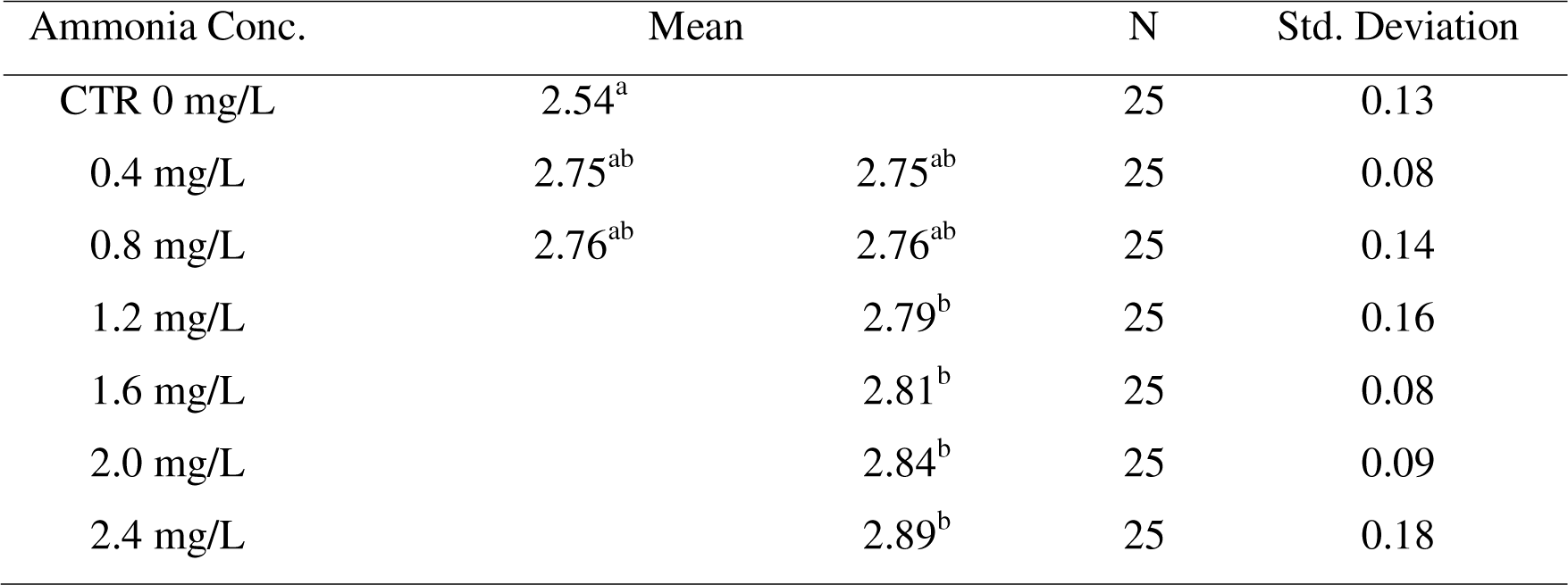
The mean scale cortisol concentrations after subjecting *O. niloticus* to ammonia treatment. The superscripts indicate significant difference at p<0.05 of the mean scale cortisol concentration following ammonia treatment. Treatment with 1.2 mg/L and above had significantly higher scale cortisol concentrations compared to the control.

**Table 6:**
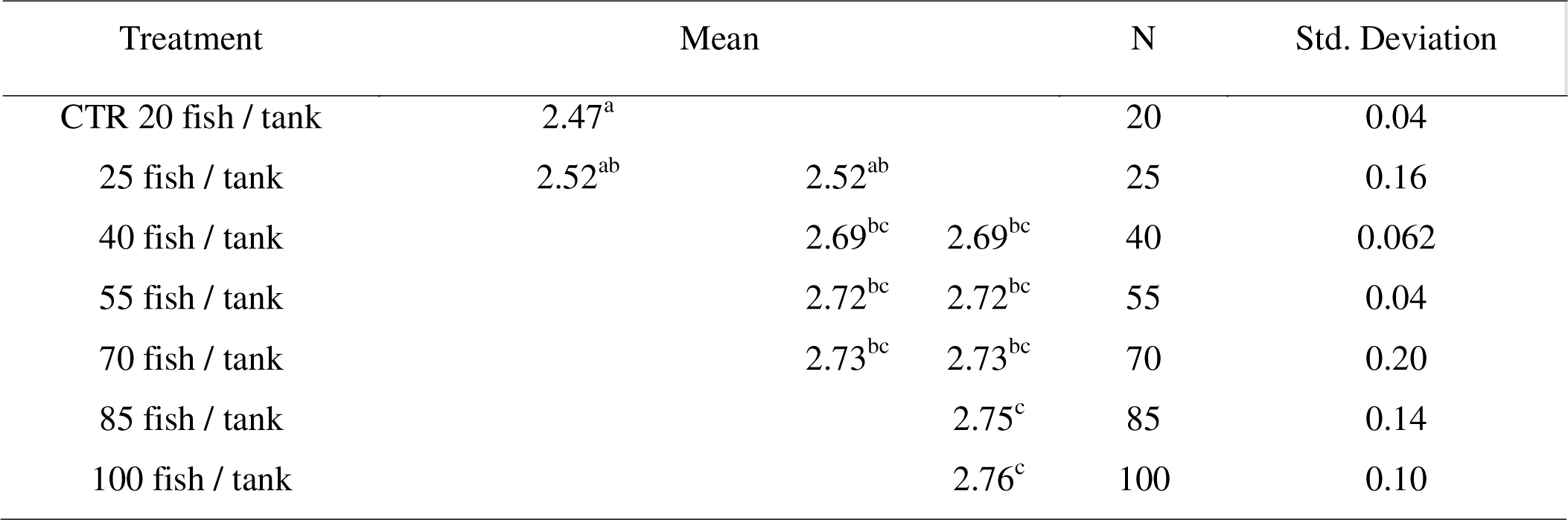
The mean scale cortisol concentrations after subjecting to stocking density treatment. The superscripts indicate significant difference at p<0.05 of the mean scale cortisol concentration following stocking density treatment. Treatment with 40 fish / tank and above had significantly higher scale cortisol concentrations compared to the control.

The ammonia treatment showed a significant difference in the levels of scale cortisol (p< 0.05) when each treatment was compared to the controls i.e., 2.54 ± 0.13 vs. 2.80 ± 0.13 mg/g for ammonia treatment (Table 5). The result clustered into two homogenous subsets between the different ammonia treatments i.e., Low ammonia concentration (0 mg/ml-0.8 mg/ml) and High ammonia concentration i.e. (1.2 mg/ml-2.4 mg/ml). In between these categories there was no significant difference (p>0.05) in the scale cortisol concentration.

The variation of scale cortisol concentration with increase in stocking density was as indicated in table 6 below. There was a positive correlation between the stocking density and the scale cortisol concentration.

### 3.4 Metabolic pathways

The effect of chronic stress on relevant metabolic pathways was evaluated by determining the differential expression of genes in the various metabolic pathways. Considering the levels of significance and the number of DEGS in the particular pathway. Mitogen activated protein kinase (MAPK) signalling pathway was the most significantly down regulated pathway in ammonia treatment [Figure 3]. Six genes in this pathway were significantly enriched following ammonia treatment (Padj ≤0.05) i.e. DUSP1 (dual specific protein phosphatase 1), NHR38 (Nuclear Hormone Receptor), HSP 72 KDa Protein 1(heat shock protein), myelocytomatosis oncogene homologue (Mych), Growth arrest and DNA damage inducible alpha a (GADD 45aa) and mitogen activated protein kinase kinase 4 (MAP2K4). Twenty-nine other genes in this pathway are also down regulated but not significantly so (Padj. ≥0.05).

**Figure 3:**
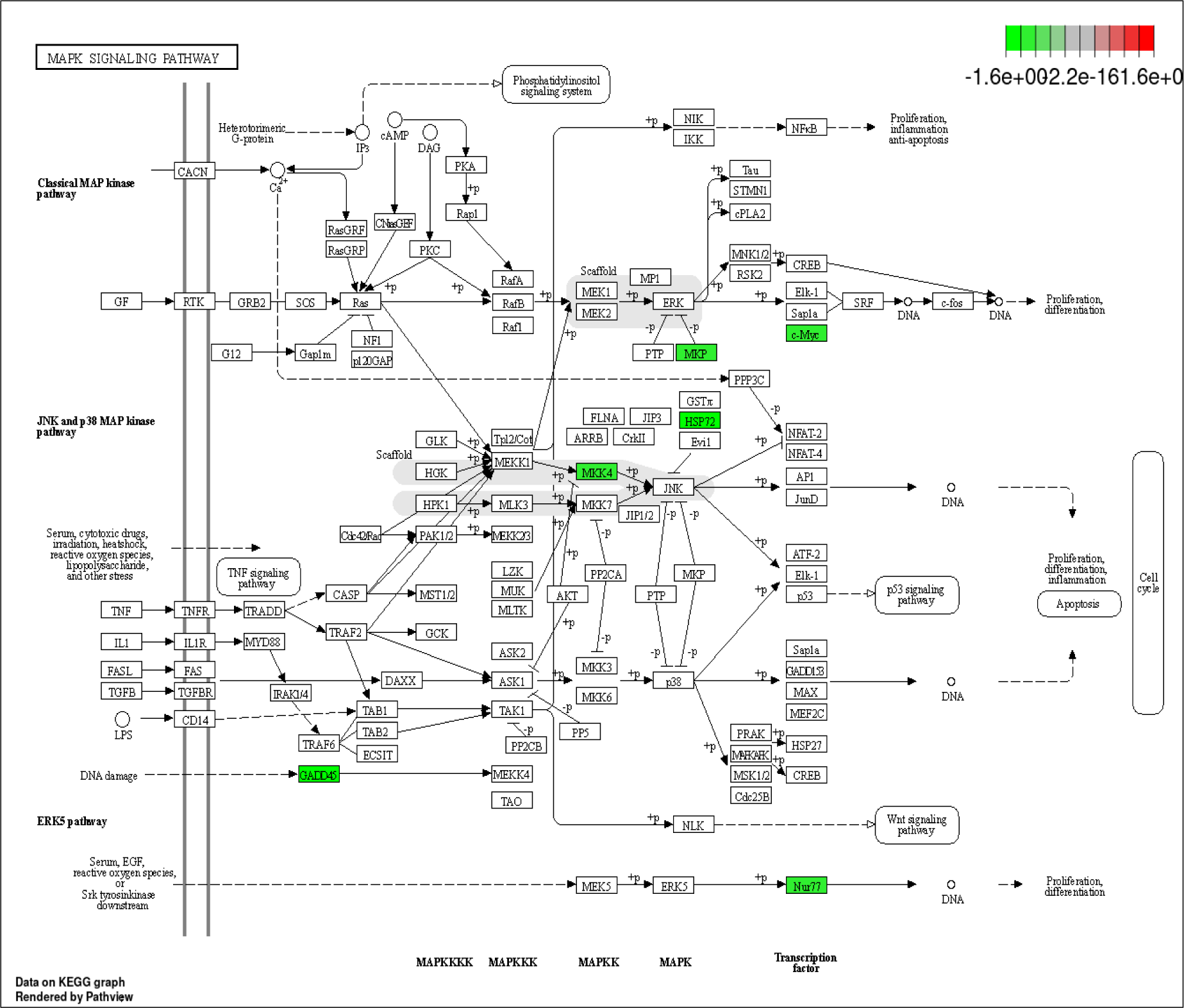
MAPK signalling pathway showing the sequence of phosphorylation leading to the activation of the respective MAPKs. The green colour indicates the significantly enriched genes under chronic stress in *O. niloticus*. The grey colour indicate the protein scaffolds that controls the phosphorylation of the MAPKs.

## Discussion

Chronic stress is the manifestation of the body’s failure to return to normal homeostatic state after stressful episodes. Stress exposes an organism to negative effects including lowered growth, reduced reproduction, and weakened immunological function. For greater productivity, disease prevention, and increased financial gain, aquaculture management practices must take stress into account and endeavour to minimize it. Glucocorticoid secretion is a typical endocrine reaction to stress that is produced in order to fuel the response to stress. In fish the major glucocorticoid produced during stress is cortisol and has been widely used as a stress indicator (Ellis et al. 2012). One of the numerous functions of cortisol in the stress response is to provide the fish with enough energy to go through the perceived stressor and subsequently resume normal activity. Cortisol plays a major role in resource reallocation during the stress. This reallocation lowers the reproductive axis, which is a prioritization of an individual’s survival over the preservation of the species. Cortisol is essential for survival during stress and controls or supports several vital metabolic, immune, and homeostatic processes. For this reason, cortisol has found wide range of application in the study of stress in many vertebrates including fish.

Under stress, fish releases cortisol as a physiological stress response. Plasma cortisol levels vary with stress levels and are indicative of the status of the culture environment (El Asey et al. 2020; Odhiambo et al. 2020). The concentration levels of cortisol are a mirror of the stress levels in the fish. Values above basal means are indicative of stressful environment. The higher plasma cortisol concentrations witnessed in the experimental fish compared to the controls were due to the independent variables introduced in the experiment I.e., ammonia and increased stocking densities and thus act as clear indicator of stress occurrence in the experimental fish.

Prolonged stress levels result in sustained high plasma cortisol levels which is eventually deposited in the scales of fish. Results of the current study indicate a positive correlation between the intensity of the stressor, the plasma cortisol concentration and the scale cortisol concentrations (Table 1, 2, 5 and 6). Scale cortisol concentration can therefore be used as an indicator of stress levels the fish has been experiencing over a period of time. This finding is in concurrence with (Aerts et al. 2015) working with common carp and concluded that scale cortisol levels could be used as long-term stress indicator. A recent study by Samaras et al. (2021), found out that isolated scale under stimulation of ACTH is able to produce cortisol on their own. Thus, fish scales besides accumulating cortisol from blood may be carrying out de novo synthesis. This coupled with the assertion that scales also participate in calcium homeostasis which would affect the cortisol storage in the scale may point to the fact that scale cortisol alone may not be sufficiently reliable in stress level determinations. This therefore points to the need for the use of a combination of markers during assessment of chronic stress in fish. An increase in fish plasma cortisol levels was accompanied by an increase Blood glucose is another important indicator of stress in fish in plasma glucose levels (Table 3 and 4). Several studies attribute glucose and cortisol levels increase with chronic stress (Hamed et al. 2012; Hamed et al. 2022; Abdel-Tawwab et al. 2023). Chronic stress brings about hyperglycaemia via enhancement of endogenous glucose synthesis and / or inhibition of glycogenesis in liver and skeletal muscles and which occurs by inhibition of glycogen synthase (Laberge et al. 2021). According to Malini et al. (2018), when fish get bigger, hyperglycaemia rises. In addition to fish size, environmental factors and the quality of the ecosystem also have an impact on hyperglycaemia.

The higher blood glucose concentration levels recorded in the current study, is an indication of the increased levels of stress the fish were exposed to. Studies by (Makaras et al. 2018; Macek et al. 2018) revealed that Plasma glucose concentrations exerts a positive correlation with increase in the level of stress. This is consistent with the findings of the current study (Table 1) where an increase in ammonia concentrations resulted in an increase in the mean plasma glucose concentrations. Similar observations were made for the stocking density stress (Table 2). Though they are the most commonly used signatures in stress level assessment, Cortisol and glucose must be combined with other measurements, such as those of other stress hormones, HSPS, blood-cell counts (preferably in long-term experiments), non-invasive techniques, and/or others, to provide a more comprehensive picture of a fish’s stress status (Zao et al. 2015). Since stress affects both physiological and biochemical processes in fish, alteration of these processes can prove a useful addition to the methods of stress determination. Most of these processes affected by stress are regulated by the levels of circulating cortisol (Yung and Giacca 2020).

The current study shows that one of the key pathways that was significantly down regulated and may have a major influence on growth performance is the Mitogen Activated Protein Kinase Signalling Pathway (MAPK). MAPK signalling pathway was significantly down regulated in ammonia treatment [Figure 3]. Six genes in this pathway were significantly enriched following ammonia treatment (Padj ≤0.05) i.e. DUSP1 (dual specific protein phosphatase 1), NHR38 (Nuclear Hormone Receptor), HSP 72 KDa Protein 1(heat shock protein), myelocytomatosis oncogene homologue (Mych), Growth arrest and DNA damage inducible alpha a (GADD 45aa) and mytogenic activated protein kinase kinase 4 (MAP2K4). Twenty-nine other genes in this pathway are also down regulated but not significantly so (Padj. ≥0.05). Chronic stress emanating from different stressors has been shown to activate the MAPK signalling pathway (Eblen 2018; Win et al. 2018). Glucocorticoids have been reported to modulate the anti-inflammatory response via the inhibition of the MAPK signalling pathway. Cortisol has specifically been shown to suppress the phosphorylation of MAPK (Dong et al. 2018), as well as the extracellular signal-regulated kinase (ERK1/2), p38MAPK and stress-activated protein kinase (JNK)/c-Jun N-terminal kinase. This suppression is mediated by DUSP-1[35] and forms the basis of the action of glucocorticoids cortisol being one of them. DUSP-1 is also referred to as (MAPK phosphatase (MKP) which mediates the de-phosphorylation of ERK (Solinas and Becattini 2016). DUSP-1 has been shown to offer a negative feedback loop for MAPK / ERK pathway (Krysan and Avila 2001). Kultz and Avila (2001) demonstrated a reduction of MAPK activity in gills of euryhaline fishes subjected to osmotic stress. Results from the current study demonstrates enrichment of MKP. This enrichment results in reduction of the phosphorylated ERK which is the effector molecule of the classical MAPK pathway. Enrichment of MKP thus down regulates ERK signalling pathway inhibiting cell differentiation and proliferation hence depressing growth. Witzel (2012), reported that the blocking of the MAPK and ERK activation also activated apoptosis.

The proto-oncogene c-MYC encodes a basic helix-loop-helix leucine zipper (bHLH-Lz) transcription factor, which plays a pivotal role in cell proliferation, metabolism, differentiation, apoptosis and tumorigenesis by transcription and activation of downstream target genes (Luscher and Larsson 1999). Marampon et al. (2006) demonstrated that the inhibition of the MEK/ERK pathway dramatically decreased c-MYC expression. Dong et al., (2019) showed that Cortisol up regulated the expression of β-catenin, c-Myc, and cyclinD1 and promoted the phosphorylation of PI3K and AKT. Cellular stress can cause the phosphorylation of c-Myc which prevents formation and binding of Myc-Max dimers to DNA. This diminishes the capacity of Myc to trigger transcription and to cause cellular proliferation, transformation, and apoptosis. P21 Acivated Kinase (Pak2) has been shown to phosphorylate Myc at the b/HLH/Z domain [56]. Results from the current study show that Nile tilapia subjected to chronic stress had significantly enriched c-Myc expression. Since c-myc enrichment diminishes cell proliferation and differentiation, this outcome is consistent with the depressed growth performance witnessed in the chronically stressed fish. The JNK MAPK pathway on the other hand is activated by two MAPK kinases (MKK4 and MKK7) that function collaboratively in response to environmental stress to optimize JNK activity. JNK is phosphorylated on Tyr residue by MKK4 and on Thr by MKK7. JNK has been demonstrated to react to a variety of cellular stress signals that are triggered by cytokines, free fatty acids, and high blood sugar. The setting and length of JNK activation have a significant impact on the outcome. While persistent JNK activation can cause cell death, transient JNK activity has the potential to promote cell growth (Yung and Giacca 2020).

Results from the current study indicate that Nile tilapia exposed to chronic stress had significantly higher MKK4 levels consistent with the higher blood glucose level witnessed in stressed fish. High glucose levels have been shown to trigger JNK activation. Phosphorylation of MKK4 leads to the activation of JNK. Prolonged activation of JNK activates apoptosis consistent with the depressed growth witnessed in this study. The coordination of gene transcription, protein synthesis, cell cycle control, apoptosis, and differentiation, orchestrated through MAPK regulation, assists in dealing with the effects of chronic stress and can therefore be viewed an important adaptation towards stress management. The transcriptional response to chronic stress and the changes in gene expression and MAPK signalling pathways in this study suggest that the aforementioned DEGs and pathways play a significant role in the mechanism behind Nile tilapia’s enhanced tolerance to chronic stress. Some of these prospective genes may be employed as candidate gene indicators of chronic stress tolerance in future breeding programs. Since the MAPK signalling pathway is essential for reacting to a variety of stressors, it is probable that manipulating the genes in this pathway will result in fish that are more resilient to the effects of climate change and more capable of producing highly.

The result of the current study increases the number of molecular markers available as tools for selective breeding in fish. Fish breeding for stress tolerance is an important tool towards increasing fish production through developing new better performing stress tolerant fish species. This will lower cost of production and minimize the occurrence of diseases and mortality in cultured fish. With the ever-increasing world population, the demand for fish and fish products continue to increase. Coupled with the dwindling catches from the wild, there is need to produce more fish in less and less area. The strain occasioned will most likely increase occurrence of stress in the cultured fish with the far-reaching consequences of depressed growth and increased mortality. As a result, this will dent the effort towards achieving freedom from hunger and malnutrition as espoused in SDG 2 as well as sink more farmers in to poverty and economic dependence contrary to SDG 1. It is therefore important to formulate policies that promote fish welfare to guard against occurrence of chronic stress in fish such as water pollution and global warming.

## Conclusion

This work demonstrates that chronic stress simultaneously increases the levels of blood glucose plasma cortisol and scale cortisol in cultured juvenile *O. niloticus*. In addition to these commonly used stress signatures, the regulation of MAP K signalling pathway can be considered as a reliable method for the assessment of chronic stress. The growth performance and welfare of cultured Nile tilapia may improve if the impact of the stressors can be reduced by providing optimal culture environments.

## Data availability statement

The primary data used to support the findings of this study are available from the corresponding author upon request.

## Authors’ contribution

JGM, designing, formal analysis and writing the original draft; POA, data curation, writing – review, editing; PO and CW, acquisition of data, analysis and interpretation of data; CW, validation, formal analysis; ETN, formal analysis, review; PO and PO, designing, conceptualization, resources and supervision. All authors read and approved the final manuscript.

## Acknowledgements

The authors would like to thank Kenya Climate Smart Agriculture Project (KCSAP) for the financial support during the study. The authors would also like to appreciate Kenya Medical Research Institute Kisumu for the technical assistance in running the ELISA tests. In addition, the authors wish to express their heartfelt gratitude to the Department of Fisheries Kakamega County for hosting the research set up. Finally, the authors would like to highly appreciate the Department of Biological Sciences, Masinde Muliro University of Science and Technology for providing valuable technical assistance and laboratory facilities throughout the research period.

## References

Abdel-Tawwab, M., Hagras, A. E., Elbaghdady, H. A. M. and Monier, M. N. (2014). Dissolved Oxygen Level and Stocking Density Effects on Growth, Feed Utilization, Physiology, and Innate Immunity of Nile Tilapia, Oreochromis niloticus. J. Appl. Aquac. 26, 340–355.

Abdel-Tawwab, M., Hamed, H. S., Monier, M. N. and Amen R. M. (2023). The ameliorative effects of dietary rosemary (*Rosmarinus officinalis*) against growth retardation, oxidative stress, and immunosuppression induced by waterborne lead toxicity in Nile tilapia fingerlings. *Ann*. Anim. Sci. 24, 139–149.

Aerts, J., Metz, J. R., Ampe B., Decostere, A., Flik G., De Saeger, S. (2015). Scales Tell a Story on the Stress History of Fish. PLOS ONE. 10(4), e0123411. 10.1371/journal.pone.0123411

Aidos, L., Cafiso, A., Serra, V., Vasconi, M., Bertotto, D., Bazzocchi, C., Radaelli, G. and Di Giancamillo A. (2020). How Different Stocking Densities Affect Growth and Stress Status of Acipenser baerii Early Stage Larvae. Anim. 10(8): 1289. 10.3390/ani10081289

Aketch, B, Angienda, P. O., Radull O. J., Waindi, E. N. (2014). Effect of stocking density on the expression of glucose transporter protein 1 and other physiological factors in the Lake Victoria Nile tilapia, Oreochromis niloticus (L.). Int. Aquat. Res. 6(2):1–8.

Balasch, J. C. and Tort, L. (2019). Netting the Stress Responses in Fish. Front. Endocrinol. 10. 10.3389/fendo.2019.00062

Cabrera-Busto, J., Mancera, J. M. and Ruiz-Jarabo, I. (2021). Cortisol and Dexamethasone Mediate Glucocorticoid Actions in the Lesser Spotted Catshark (*Scyliorhinus canicula*). Biol 11(1), 56. 10.3390/biology11010056

Carbajal, A., Monclús, L., Tallo-Parra, O., Sabes-Alsina, M., Vinyoles, D. and Lopez-Bejar, M. (2018). Cortisol detection in fish scales by enzyme immunoassay: Biochemical and methodological validation. J. Appl. Ichthyol. 34(4): 967–970. 10.1111/jai.13674

Cargnello, M. and Roux, P. P. (2011). Activation and Function of the MAPKs and Their Substrates, the MAPK-Activated Protein Kinases. Microbiol. Mol. Biol. Rev. 75(1), 50–83. 10.1128/mmbr.00031-10

Dong, J., Li, J., Li, J., Cui, L., Meng, X., Qu, Y., Wang, H. (2019). The proliferative effect of cortisol on bovine endometrial epithelial cells. Reprod. Biol. Endocrinol. 17(1). 10.1186/s12958-019-0544-1

Dong, J., Qu, Y., Li, J., Cui, L., Wang, Y., Lin, J. and Wang, H. (2018). Cortisol inhibits NF-κB and MAPK pathways in LPS activated bovine endometrial epithelial cells. Int Immunopharmacol. 56: 71–77. 10.1016/j.intimp.2018.01.021

Eanes, L. and Patel, Y. M. (2016). Inhibition of the MAPK pathway alone is insufficient to account for all of the cytotoxic effects of naringenin in MCF-7 breast cancer cells. Biochimie Open. 3: 64–71. 10.1016/j.biopen.2016.09.004

Eblen, S. T. (2018). Extracellular-Regulated Kinases: Signalling From Ras to ERK Substrates to Control Biological Outcomes. Adv. Cancer Res. 138, 99–142. 10.1016/bs.acr.2018.02.004

El Asey, M. A., Reda R. M., Salah A. S., Mahmoud, M. A. and Dawood, M. A. O. (2020). Overall performances of Nile tilapia (*Oreochromis niloticus*) associated with using vegetable oil sources under suboptimal temperature. Aquac. Nutr. 26(4), 1154–1163.

Ellis T., Yildiz H. Y., López-Olmeda, J., Spedicato, M. T., Tort, L., Øverli, Ø. and Martinis M. I. C. (2012). Cortisol and finfish welfare. Fish. Physiol. Biochem. 38, 163–188. doi: 10.1007/978-94-007-5383-9_11

Faught, E., Vijayan, M. M. (2016). Mechanisms of cortisol action in fish hepatocytes. Comparative Biochemistry and Physiology Part B: *Biochem*. Mol. Biol. 199, 136–145. 10.1016/j.cbpb.2016.06.012

Food and Agriculture Organization of the United Nations (2022) *The State of World Fisheries and Aquaculture-Towards blue transformation*; Rome, Italy.

Goda, M., Shaheen, A. A. M. and Hamed, H. S. (2023). Potential role of dietary parsley and/or parsley nanoparticles against zinc oxide nanoparticles toxicity induced physiological, and histological alterations in Nile tilapia, Oreochromis niloticus. Aquac. Rep. 28(101425).

Godoy-Olmos, S., Jauralde, I., Monge-Ortiz, R., Milián-Sorribes M. C., Jover-Cerdá, M,. Tomás-Vidal, A. and Martínez-Llorens, S. (2022). Influence of diet and feeding strategy on the performance of nitrifying trickling filter, oxygen consumption and ammonia excretion of gilthead sea bream (Sparus aurata) raised in recirculating aquaculture systems. Aquac Int. 30(2): 581–606. 10.1007/s10499-021-00821-3

Hamed, H. S. and Abdel-Tawwab, M. (2021). Dietary pomegranate (*Punica granatum*) peel mitigated the adverse effects of silver nanoparticles on the performance, haemato-biochemical, antioxidant, and immune responses of Nile tilapia fingerlings. Aquac. 540(7), 736–742.

Hamed, H. S., Ali, R. M., Shaheen, A. A. and Hussein, N. M. (2021). Chitosan nanoparticles alleviated endocrine disruption, oxidative damage, and genotoxicity of Bisphenol-A-intoxicated female African cat fish. Comp. Biochem. Physiol. Part (C). 248.

Hamed, H. S., Ismal, S. M. and Abdel-Tawwab M. (2022). Modulatory effects of cinnamon (*Cinnamomum zeylanicum*) against waterborne lead toxicity in Nile tilapia fingerlings: Growth performance, haemato-biochemical, innate immunity and hepatic antioxidant capacity. Aquac Rep. 25 (101190).

Hamed, H. S., Amen, R. M., Elelemi, A. H, Mahboub H. H., Elabd, H,. Abdelfattah, M. A,. Abdel-Moniem, H., El-Beltagy, M. A., Alkafafy, M., Yassin, E. M. M. and Ismail, A. K. (2022). Effect of dietary Moringa oleifera leaves nanoparticles on Growth performance, physiological, immunological responses, and liver antioxidant biomarkers in Nile tilapia (*Oreochromis niloticus*) against Zinc oxide nanoparticles toxicity. Fishes. 7 (360).

Harper, C. and Wolf, J. C.(2009). Morphologic Effects of the Stress Response in Fish. ILAR J 50(4), 387–396. 10.1093/ilar.50.4.387

Hoppstädter, J. and Ammit, A. J. (2019). Role of Dual-Specificity Phosphatase 1 in Glucocorticoid-Driven Anti-inflammatory Responses. Front Immunol. 10. 10.3389/fimmu.2019.01446

Jensen, J., Rustad, P. I., Kolnes, A. J., and Lai, Y. C. (2011). The role of skeletal muscle glycogen breakdown for regulation of insulin sensitivity by exercise. Front Physiol. 2,112. 10.3389/fphys.2011.00112

Jia, R,. Wang, L., Hou, Y., Feng, W., Li, B. and Zhu, J. (2022) Effects of Stocking Density on the Growth Performance, Physiological Parameters, Redox Status and Lipid Metabolism of Micropterus salmoides in Integrated Rice–Fish Farming Systems. Antioxidants. 11(7), 1215 10.3390/antiox11071215

Kassel, O. (2001). Glucocorticoids inhibit MAP kinase via increased expression and decreased degradation of MKP-1. The EMBO J. 20(24), 7108–7116. 10.1093/emboj/20.24.7108

Kennedy, E. K. C. and Janz, D. M. (2022). First Look into the Use of Fish Scales as a Medium for Multi-Hormone Stress Analyses. Fishes. 7(4), 145 10.3390/fishes7040145

Krysan, P. J. and Colcombet, J. (2018). Cellular Complexity in MAPK Signalling in Plants: Questions and Emerging Tools to Answer Them. Front Plant Sci. 9. 10.3389/fpls.2018.01674

Kültz, D. and Avila, K. (2001). Mitogen-activated protein kinases are in vivo transducers of osmosensory signals in fish gill cells. Comp. Biochem. Physiol. Part B: Biochem Mol Biol. 129(4), 821–829 10.1016/s1096-4959(01)00395-5

Kyriakis, J. M. and Avruch, J. (2012). Mammalian MAPK Signal Transduction Pathways Activated by Stress and Inflammation: A 10-Year Update. Physiol Rev. 92(2), 689–737 10.1152/physrev.00028.2011

Laberge, F., Yin-Liao, I. and Bernier, N. J. (2019). Temporal profiles of cortisol accumulation and clearance support scale cortisol content as an indicator of chronic stress in fish. Conserv Physiol. 7(1). 10.1093/conphys/coz052

Lai, F., Royan, M. R., Gomes, A. S., Espe, M., Aksnes, A., Norberg, B., Gelebart, V. and Rønnestad I. (2021). The stress response in Atlantic salmon (Salmo salar L.): identification and functional characterization of the corticotropin-releasing factor (crf) paralogs. Gen Comp Endocrinol. 313, 113894. 10.1016/j.ygcen.2021.113894

Lüscher, B. and Larsson, L. G. (1999). The basic region/helix[–[loop[–[helix/leucine zipper domain of Myc proto-oncoproteins: Function and regulation. Oncogene. 18(19), 2955–2966. 10.1038/sj.onc.1202750

Macek, P., Cliff, M. J., Embrey, K. J., Holdgate, G. A., Nissink, J. W. M., Panova, S., Waltho, J. P. and Davies, R. A. (2018) Myc phosphorylation in its basic helix-loop-helix region destabilizes transient α-helical structures, disrupting Max and DNA binding. The J Biol Chem. 293(24), 9301– 9310. 10.1074/jbc.RA118.002709

Mahboub, H. H., Amen, R. M., El-Beltagy, M. A,. Ramah, A., Abdelfattah, A. M., El-Beltagi, H. S., Shalaby, T. A., Ghazzawy, H. S., Ramadan, K. M. A., Alhajji, A. H. M. and Hamed, H. S. (2022) Ameliorative effect of Quercetin against abamectin induced hemato-biochemical alterations and hepatorenal oxidative damage in Nile tilapia, Oreochromis niloticus. Anim. 12(23), 3429:1-16

Makaras, T., Razumienė, J,. Gurevičienė, V., Šakinytė, I., Stankevičiūtė, M. and Kazlauskienė, N. (2020). A new approach of stress evaluation in fish using β-d-Glucose measurement in fish holding-water. Ecol Indic. 109, 105829. 10.1016/j.ecolind.2019.105829

Malini, D. M,. Madihah, M., Apriliandri, A. F. and Arista, S. (2018). Increased Blood Glucose Level on Pelagic Fish as Response to Environmental Disturbances at East Coast Pangandaran, West Java. IOP Conference Series: Earth Environ Sci. 166, 012011. 10.1088/1755-1315/166/1/012011

Mansour AT, Amen RM, Mahboub HH, Shawky SM, Orabi SH, Ramah A, Hamed HS (2023) Exposure to oxyfluorfen-induced hematobiochemical alterations, oxidative stress, genotoxicity, and disruption of sex hormones in male African catfish and the potential to confront by Chlorella vulgaris. *Comp. Biochem. Physiol. Part (C)*, Toxicol Pharmacol. 267.

Marampon, F., Ciccarelli, C., Zani, B. M. (2006). Down-regulation of c-Myc following MEK/ERK inhibition halts the expression of malignant phenotype in rhabdomyosarcoma and in non muscle-derived human tumors. Mol Cancer. 5(1). 10.1186/1476-4598-5-31

Martos-Sitcha, J. A., Mancera, J. M., Prunet, P. Magnoni, L. J. (2020). Editorial: Welfare and Stressors in Fish: Challenges Facing Aquaculture. Front Physiol. 11. 10.3389/fphys.2020.00162

Montero, D., Marrero, M., Izquierdo, M., Robaina, L., Vergara, J. and Tort, L. (1999). Effect of vitamin E and C dietary supplementation on some immune parameters of gilthead seabream (S*parus aurata*) juveniles subjected to crowding stress. Aquac. 171(3–4), 269–278. 10.1016/s0044-8486(98)00387-1

Odhiambo, E., Angienda P. O., Okoth, P., Onyango, D. (2020). Stocking Density Induced Stress on Plasma Cortisol and Whole Blood Glucose Concentration in Nile Tilapia Fish (*Oreochromis niloticus*) of Lake Victoria, Kenya. Int J Zool. 1–8. 10.1155/2020/9395268

Oké, V. and Goosen, N. J. (2019). Corrigendum to ‘The effect of stocking density on profitability of African catfish (*Clarias gariepinus*) culture in extensive pond systems’. Aquac. 507, 385–392. 10.1016/j.aquaculture.2019.734270

Pasnik, D. J., Evans, J. J. and Klesius, P. H. (2008). Influnce of tricanemethanesulfonate on *Streptococcus agalactiae* vaccination of Nile tilapia (*Oreochromis niloticus*). J Vet Res. 2(2), 28– 33

Pickering A. D. and Pottinger, T. G. (1989). Stress responses and disease resistance in salmonid fish: Effects of chronic elevation of plasma cortisol. Fish Physiol Biochem. 7(1–6), 253–258. 10.1007/bf00004714

Rodriguez-Barreto, D., Rey, O., Uren-Webster, T. M., Castaldo, G., Consuegra, S. and Garcia de Leaniz C. (2019) Transcriptomic response to aquaculture intensification in Nile tilapia. Evol Appl. 12(9), 1757–1771. 10.1111/eva.12830

Roque d’orbcastel, E., Bettarel, Y., Dellinger, M., Sadoul, B., Bouvier, T., Amandé, J. M., Dagorn, L. and Geffroy, B. (2021). Measuring cortisol in fish scales to study stress in wild tropical tuna. Environ. Biol. Fishes. 104(6), 725–732. 10.1007/s10641-021-01107-6

Sadoul, B. and Geffroy, B. (2019) Measuring cortisol, the major stress hormone in fishes. J. Fish Biol. 94(4), 540–555. 10.1111/jfb.13904

Samaras, A., Dimitroglou, A., Kollias, S., Skouradakis, G., Papadakis, I. E. and Pavlidis, M. (2021). Cortisol concentration in scales is a valid indicator for the assessment of chronic stress in European sea bass, Dicentrarchus labrax L. Aquac 545: 737257. 10.1016/j.aquaculture.2021.737257

Sapolsky, R. M., Romero, L. M. and Munck, A. U. (2000). How Do Glucocorticoids Influence Stress Responses? Integrating Permissive, Suppressive, Stimulatory, and Preparative Actions. Endocr. Rev. 21(1), 55–89. 10.1210/edrv.21.1.0389

Solinas, G. and Becattini, B. (2016). JNK at the crossroad of obesity, insulin resistance, and cell stress response. Mol. Met. 6(2), 174–184. 10.1016/j.molmet.2016.12.001

Thorarensen, H., Kubiriza, G. and Imsland. A. (2015). Experimental design and statistical analyses of fish growth studies. Aquac. 448, 483–490.

Tohamy, H. G., Lebda, M. A., Sadek, K. M., Elfeky, M. S., El-sayed, Y. S., Samak, D. H., Hamed, H. S. and Abouzed, T. K. (2022). Biochemical, molecular and cytological impacts of alpha-lipoic acid and Ginkgo biloba in ameliorating testicular dysfunctions induced by silver nanoparticles in rats. Environ. Sci. Pull. Res. Int. 29(250, 38198–38211.

Vercauteren, M., Van, Hoey, G., Decostere, A., Boyen, F., Ampe, B., Devriese, L. and Chiers, K. (2021). Influence of Pathogens, Fish-Related Characteristics, and Environmental Factors on the Development of Skin Ulcerations in Wild Common Dab (*Limanda limanda*) From the North Sea. J. Wildl. Dis. 57(2). 10.7589/Jwd-D-20-00088

Volpato, G. and Barreto, R. (2001). Environmental blue light prevents stress in the fish Nile tilapia. Braz. J. Med. Biol. Res. 34(8), 1041–1045. 10.1590/s0100-879x2001000800011

Wang, J., Zhou, J. Y., Kho, D., Reiners, J. J. and Wu, G. S. (2016). Role for DUSP1 (dual-specificity protein phosphatase 1) in the regulation of autophagy. Autophagy. 12(10), 1791–1803. 10.1080/15548627.2016.1203483

Win, S., Than, T. A. and Kaplowitz, N. (2018). The regulation of JNK Signalling pathways in cell death through the interplay with mitochondrial SAB and Upstream Post-Translational Effects. Int. J. Mol. Sci. 19(11), 3657. 10.3390/ijms19113657

Witzel, F., Maddison, L. and Blüthgen, N. (2012). How scaffolds shape MAPK signalling: what we know and opportunities for systems approaches. Front. Physiol. 3. 10.3389/fphys.2012.00475

Yung, J. H. M. and Giacca, A. (2020). Role of c-Jun N-terminal Kinase (JNK) in Obesity and Type 2 Diabetes. Cells. 9(3), 706. 10.3390/cells9030706

Zeitoun, M. M., EL-Azrak, K. E. D. M., Zaki, M. A., Nemat-Allah, B. R. and Mehana, E. S. E. (2016). Effects of ammonia toxicity on growth performance, cortisol, glucose and hematological response of Nile Tilapia (Oreochromis niloticus). *Aceh*. J. Anim. Sci. 1(1), 21–28. 10.13170/ajas.1.1.4077

Zenth, F., Corlatti, L., Giacomelli, S., Saleri, R., Cavalli, V., Andrani, M., Donini, V. (2022). Hair cortisol concentration as a marker of long-term stress: sex and body temperature are major determinants in wild-living Alpine marmots. Mamm. Biol. 102, 2083–2089. 10.1007/s42991-022-00264-0

Zhao, Q., Assimopoulou, A. N., Klauck, S. M., Damianakos, H., Chinou, I., Kretschmer, N., Rios, J.L., Papageorgiou, V. P., Bauer, R., Efferth, T. (2015). Inhibition of c-MYC with involvement of ERK/JNK/MAPK and AKT pathways as a novel mechanism for shikonin and its derivatives in killing leukemia cells. Oncotarget. 6(36), 38934–38951. 10.18632/oncotarget.5380

Zhu, C. D., Wang, Z. H. and Yan, B. (2013). Strategies for hypoxia adaptation in fish species: a review. J. Comp. Physiol. B. 183(8), 1005–1013. 10.1007/s00360-013-0762-3

